# Programmable antigen-presenting scaffolds expand rare tumor-antigen-specific T cells from peripheral blood

**DOI:** 10.64898/2026.10.05.756899

**Authors:** Jessica DeVilla, Yoav Binenbaum, Nicoletta Cieri, David Zhang, Helen Ji, Vittoria Martinolli D’Arcy, Giacomo Oliveira, Michael Manos, Donna Neuberg, Jerome Ritz, Patrick A. Ott, David Mooney, Catherine J. Wu

## Abstract

Adoptive cellular therapies can be effective against solid tumors but rely on the identification and expansion of rare tumor-reactive T cells (TRTs). Here, we utilize antigen-presenting cell-mimetic scaffolds (APC-ms), a modular biomaterial platform that presents defined peptide-HLA and co-stimulatory ligands on a fluid lipid bilayer to selectively recover rare antigen-specific T cells from peripheral blood. Independent variation of peptide-HLA, co-stimulatory ligands, and cytokine support defined a formulation that enriched MART-1-specific T cells to above 2% of CD8^+^ T cells in 35 of 50 HLA-A*02:01^+^ healthy donors from low or undetectable baseline frequencies. Compared with mo-DCs, APC-ms achieved greater fold expansion of MART-1-specific T cells with increased functional avidity and preservation of an early differentiation phenotype, and generated a higher number of MART-1-specific clonotypes. In five of eight patients with treatment-naïve melanoma, APC-ms could enrich MART-1-specific T cells to a median of 13.7% of CD8^+^ cells (range 0.30 – 57.2%) Moreover, APC-ms could expand T cells specific for an HLA-A*24:01-restricted personal neoantigen that were previously undetectable in peripheral blood of a patient with melanoma. These findings establish APC-ms as a tunable platform for the recovery of rare TRTs from peripheral blood, supporting its further development for therapeutic TCR discovery and personalized adoptive cell therapy.

## Main

Adoptive cellular therapies, including tumor-infiltrating lymphocyte (TIL) therapy and T cell receptor (TCR)-engineered T cell therapy (TCR-T), are expanding the therapeutic landscape for solid tumors^1–3^. TIL therapy relies on the recovery and expansion of tumor-reactive T cells (TRTs) from resected lesions, whereas TCR-T requires genetic transfer of a previously identified tumor-reactive TCR. A critical challenge with both is the identification and expansion of rare tumor-reactive populations in individual patients, given the known high abundance of bystander clonotypes in TIL and the relative rarity of tumor-reactive populations at the tumor site^4^. Moreover, TRTs found in TIL are often characterized by phenotypes of chronic antigen stimulation and exhaustion which limit both their capacity for tumor reactivity and their intrinsic expansion potential^5–8^.

Peripheral blood offers a relatively untapped, minimally invasive source of native TRTs, including clonotypes that may be absent or poorly represented within an individual tumor specimen. In contrast to TIL, circulating TRTs have the advantage that they can retain markers of functional fitness and display memory phenotypes that may favor ex vivo expansion^9^. Indeed, several efforts have identified tumor-reactive TCRs from peripheral blood, including those recognizing tumor-associated antigens (TAAs) and patient-specific neoantigens^8,10–12^. However, neoantigen-reactive T cells are present at extremely low precursor frequencies, estimated to range from 0.02-0.0007% of the peripheral blood repertoire^8,10^. Efforts to enrich these populations have relied on sorting for markers of exhaustion such as PD-1 and CD39^10,12^. However, the rarity of these TRTs remains a fundamental barrier to their recovery.

Existing methods for *in vitro* enrichment of TRTs rely on antigen-presenting cells (APCs), either patient-derived or engineered, to provide antigen-specific priming and stimulation. Peptide-pulsed monocyte-derived dendritic cells (mo-DCs) remain the standard for antigen-specific expansion, but require labor-intensive and patient-specific manufacturing. Donor variability in viability and phenotype compromise recovery of low-frequency antigen-specific clonotypes, particularly among patients in which DCs may display impaired antigen-presentation and priming of T cells^13,14^. This limitation is consequential when attempting to recover rare TRTs that may require narrowly defined stimulation conditions because of prior antigen exposure or tumor-induced tolerance. Artificial APCs (aAPCs) offer an attractive solution, circumventing the need for patient-derived APCs while allowing independent control over presentation of stimulatory molecules such as peptide-HLA (pHLA) molecules, co-stimulatory ligands, and cytokines. In recent years, a range of aAPC designs have been explored to selectively expand antigen-specific T cells, including polymeric scaffolds^15–17^, microparticles^18–20^, and hydrogels^21^, illustrating the feasibility of targeted antigen-specific expansion with aAPCs.

Here we utilize an aAPC composed of high-aspect-ratio mesoporous silica rods (MSRs) coated with a supported lipid bilayer and functionalized with biotinylated ligands through a streptavidin intermediate^22,23^ to selectively expand TRTs from peripheral blood. This system, termed APC-mimetic scaffolds (APC-ms), allows for presentation of stimulatory ligands with lateral mobility along the bilayer to mimic the dynamic interactions between T cells and cellular APCs. The platform also offers modularity for inclusion of pHLA and costimulatory molecules while allowing for precise tuning of ligand density and stoichiometry. In prior studies, APC-ms presenting αCD3/αCD28 expanded primary human T cells from peripheral blood two-to ten-fold more efficiently than commercial beads,^23^ tuned CAR-T cell phenotype and function with varying ligand-density,^24^ and expanded Epstein-Barr virus-specific T cells capable of recognizing cognate peptide-pulsed targets^23^. Herein, we leverage APC-ms as a controlled biomaterial interface to define conditions for expanding rare, antigen-specific T cells from peripheral blood. We then demonstrate that APC-ms outperform mo-DCs in enriching antigen-specific T cell clonotypes. Finally, we demonstrate that APC-ms can expand rare, patient-derived T cell populations specific to both a shared tumor-associated antigen and a validated personal neoantigen.

### Tuning APC-ms to expand MART-1-specific T cells from blood

To independently control the three canonical signals of T cell activation (peptide-HLA presentation, co-stimulation, and cytokine support), APC-ms were applied to define conditions that selectively recover and expand antigen-specific T cells from the peripheral blood in healthy donors (Fig. 1a, Extended Data Fig. 1). MART-1, the melanocyte differentiation marker and TAA in melanoma, was used as a model antigen due to its known large precursor frequency in HLA-A*02:01 individuals (greater than one in 2,500 CD8+ T cells)^25^. Purified T cells from HLA-A*02:01^+^ healthy donors were co-cultured with APC-ms presenting the anchor-modified MART-126-35(A27L) epitope ELAGIGILTV on HLA-A*02:01, together with defined co-stimulatory ligands and cytokine regiments^26,27^. Antigen-specific CD8^+^ T cells were quantified on day 12 using pHLA multimers and evaluated by MART-1-specific frequency among CD8^+^ T cells, MART-1-specific fold expansion from baseline, absolute antigen-specific yield and the proportion of MART-1-specific cells retaining a CD45RA+CD62L+ stem cell memory-associated phenotype (T_SCM_) (Fig. 1b, Extended Data Fig. 2).

**Figure 1:**
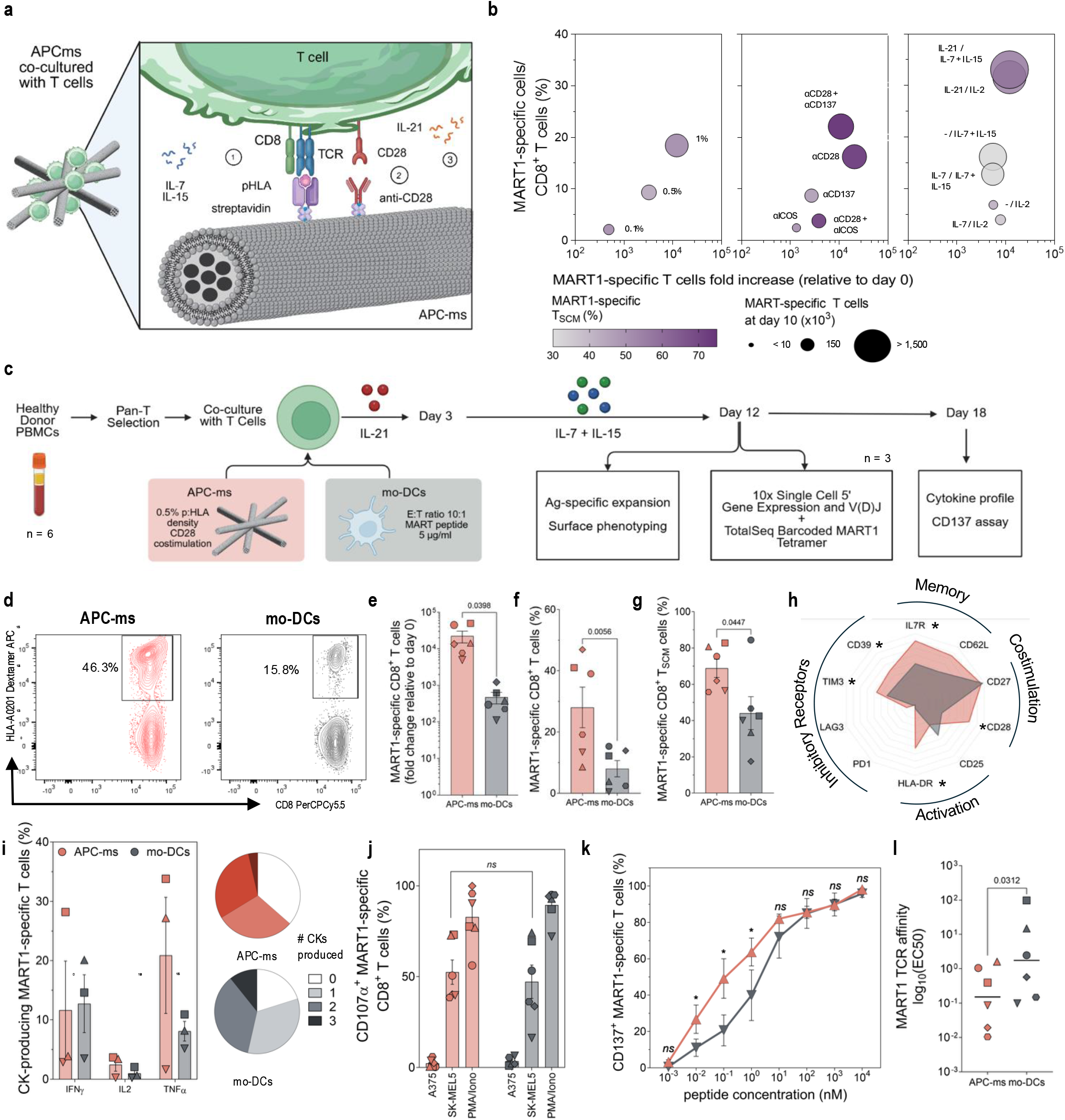
Optimization of APC-ms for expansion of MART-1-specific T cells from peripheral blood and comparison with mo-DCs. **(a)** Graphic of APC-ms co-culture with T cells. APC-ms present peptide-HLA for TCR stimulation and co-stimulatory ligands (αCD28 shown) (created with BioRender.com). Cytokine are supplemented in the culture medium. **(b)** APC-ms formulations were optimized for MART-1-specific T cell expansion by varying particle presentation density, costimulatory molecules, and cytokine support. T cells were isolated from peripheral blood of healthy donors and co-cultured with APC-ms for 12 days. MART-1-specific T cell enrichment (dextramer^+^ cells among CD8^+^ T cells), fold expansion, frequency of MART-1-specific T cells within the T_SCM_ compartment (CD62L^+^ CD45RA^+)^, and absolute MART-1-specific T cell numbers are shown. Data represent the mean across five healthy donors. In the right-most panel, cytokines are represented as: days 0-2 of expansion/day 3+ of expansion. Hyphens signify no cytokine was added. **(c)** Experimental schematic for comparison of APC-ms and mo-DC-mediated expansion. T cells from six heathy donors were expanded in triplicate. **(d)** Representative flow cytometry plots showing MART-1-specific (dextramer^+^) CD8^+^ T cells in a single donor expanded with APC-ms or mo-DCs. **(e)** MART-1-specific expansion, **(f)** MART-1-specific CD8^+^ T cell frequency, and **(g)** MART-1-specific T_SCM_ frequency. Each symbol represents a different donor. **(h)** Radar plot showing the mean expression of T cell phenotypic markers across 6 donors after expansion with APC-ms or mo-DCs. **(i)** Intracellular flow cytometry analysis of IFNγ, IL-2, and TNFα production following coculture with MART-1^+^ HLA-A*02:01^+^ SK-MEL-5 cells. Pie charts represent the proportions of MART-1-specific T cells that did not produce cytokine, produced only a single cytokine, or produced several of the aforementioned cytokines (n = 3 donors). **(j)** CD107α degranulation of APC-ms- and mo-DC-expanded T cells following co-culture with MART-1^+^ HLA-A*02-01^+^ SK-MEL-5 cells, MART-1^-^ HLA-A*02-01^+^ A375 cells, or stimulation with PMA/Ionomycin (n=5 donors). **(k)** Normalized CD137 expression following 20 hours of coculture with peptide-pulsed T2 cells and comparison of functional avidity (EC50 of CD137 curve) for T cells expanded with APC-ms or mo-DCs (n = 6 donors). Error bars represent mean ± SEM. P values were calculated using donor-level paired t tests.

Increasing the density of pHLA and αCD28 displayed on aAPC-ms enhanced MART-1-specific enrichment and total expansion (Fig. 1b, Extended Data Fig. 3). Among the co-stimulatory ligand combinations tested (αCD28, αCD137, or αICOS), αCD28 co-stimulation alone was necessary and sufficient to achieve robust MART-1-specific expansion (Fig. 1b, Extended Data Fig. 3). Finally, sequential exposure to IL-21 (on days 0-3) followed by low-dose IL-7 and IL-15 produced the greatest MART-1-specific expansion while retaining the largest T_SCM_ fraction, consistent with established roles for these cytokines in CD8^+^ T cell priming and memory maintenance (Fig. 1b, Extended Data Fig. 3)^28–30^. Supplementation with IL-21 during the first three days of culture increased antigen-specific enrichment and resulted in a less differentiated phenotype (Fig. 1b). Based on these results, we completed subsequent experiments using a tuned reference formulation consisting of 0.5 mol% lipid biotinylation, αCD28 co-stimulation, and sequential IL-21 followed by IL-7 and IL-15.

### APC-ms increase MART-1-specific expansion and functional avidity relative to mo-DCs while preserving functionality

APC-ms were compared with autologous MART-1 peptide-pulsed mo-DCs generated from six healthy donors. T cells from each donor were expanded in parallel with APC-ms or mo-DCs (Fig. 1c, Extended Data Fig. 4a-d). In the most responsive donor, MART-1-specific cells reached 46.3% of CD8^+^ T cells after APC-ms expansion compared with 15.8% after mo-DC expansion (Fig. 1d). By day 12, APC-ms generated a greater MART-1-specific CD8^+^ T cell frequency (p = 0.006) and greater fold expansion from baseline (p = 0.039) than mo-DCs (Fig. 1e,f, Extended Data Fig. 4f). APC-ms-expanded MART-1-specific cells contained a larger CD45RA^+^CD62L^+^ (T_SCM_) compartment than mo-DC-expanded cells (p = 0.045, Fig. 1g) and showed higher surface expression of IL-7Rα, HLA-DR, CD28, CD39, and TIM-3 (Fig. 1h, Extended Data Fig. 4f). This phenotype combines IL-7Rα expression with markers of activation, consistent with an *ex vivo* expanded population that retains features associated with T cell fitness. Both APC-ms- and mo-DC expanded MART-1-specific cells produced interferon-γ (IFNγ), tumor necrosis factor-α (TNFα) and IL-2 after challenge with MART-1^+^ HLA-A*02:01^+^ SK-MEL-5 cells, with polyfunctional cytokine production observed in the three donors tested (Fig. 1i). CD107α degranulation was comparable between expansion platforms in six donors and was low against MART-1^-^ HLA-A*02:01^+^ A375 cells relative to SK-MEL-5 cells, supporting antigen-dependent effector function (Fig. 1j). Because peptide responsiveness is a desirable feature of TRTs, peptide dose-response curves for CD137 upregulation were compared following expansion with APC-ms or monocyte-derived DCs (Fig. 1k). APC-ms selectively expanded T cells with greater functional avidity compared to mo-DCs, as reflected by 38-fold lower EC50 (APC-ms: 0.5nM, mo-DCs: 19.2nM, p=0.031) (Fig. 1l).

MART-1-specific expansion with APC-ms was also compared with Xynapse, a commercial dextran-based antigen-presenting scaffold (Immudex)^15^. APC-ms produced greater MART-1-specific enrichment and fold expansion across three donors (Extended Data Fig. 5), providing an additional benchmark against an alternative synthetic antigen-presenting platform.

### APC-ms expanded T cells demonstrate clonal expansion and signatures of cytotoxicity and memory

To further assess differences between APC-ms and mo-DC expanded T cells, we performed 5′ single-cell RNA sequencing with paired V(D)J profiling and CITE-seq on day-12 cultures from three donors expanded in parallel with APC-ms or mo-DCs. DNA-barcoded MART-1 and control tetramers were used to distinguish antigen-specific T cells from non-specific background (Extended Data Fig. 6, Supplementary Table 1). Unsupervised clustering resolved MART-1-specific T cells into three distinct clusters (Fig. 2a). MART-1-specific cells from APC-ms were enriched in clusters 1 and 2, while those from mo-DCs were enriched in cluster 0, illustrating a divergence in cell state (Extended Data Fig. 7). APC-ms-expanded-MART-1-specific T cells displayed significantly higher expression of markers associated with early differentiation and T cell fitness (*SELL, IL7R*) and of cytotoxic potential (*GZMB, IFNG, ZNF683*), consistent with ex vivo activation peak status (Fig. 2b, Supplementary Table 2).

**Figure 2:**
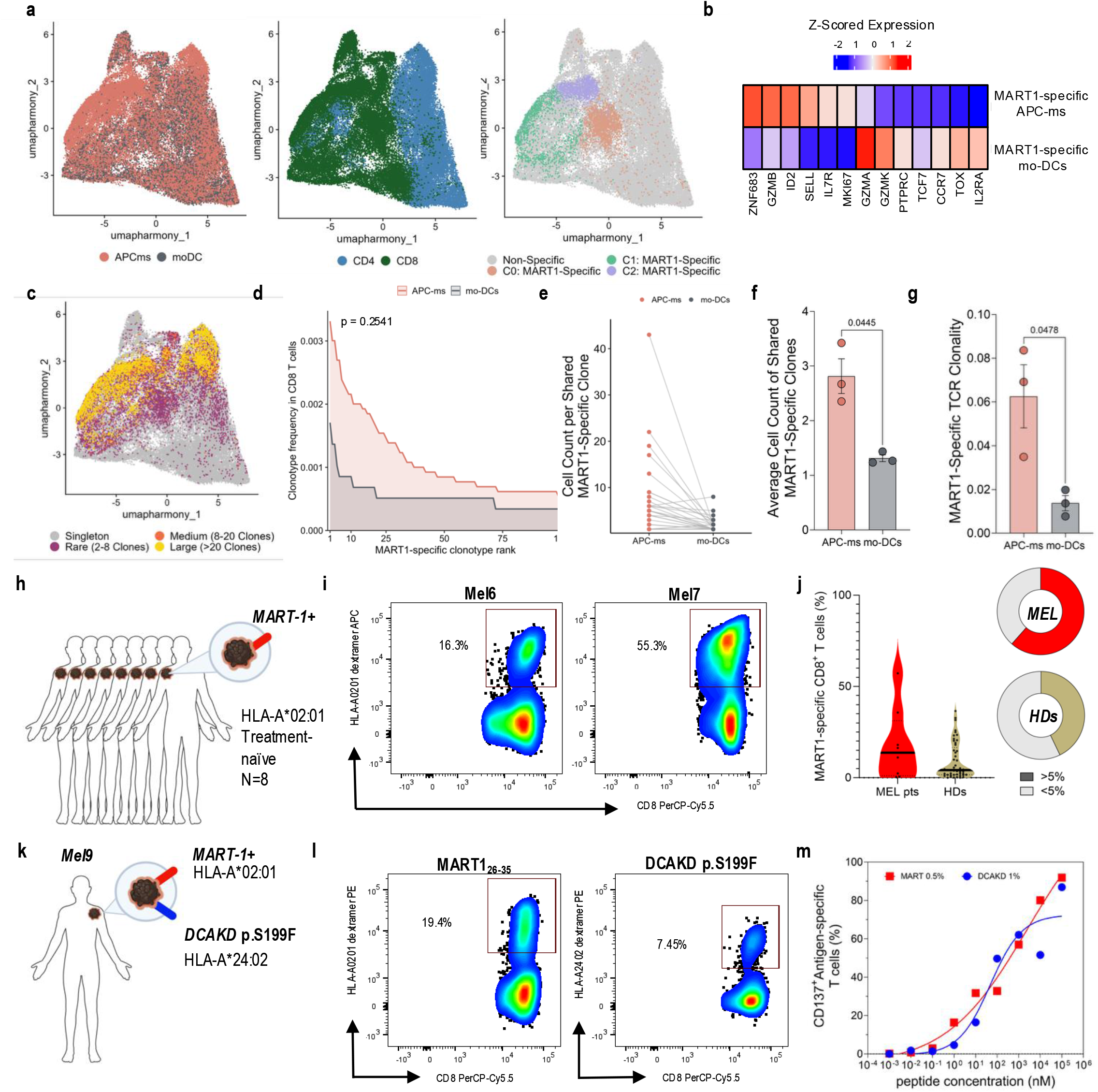
Transcriptomic characterization of expanded MART-1-specific T cells and antigen-specific expansion from peripheral blood of melanoma patients. T cells from 3 healthy donors were co-cultured with APC-ms and mo-DCs in parallel and labeled with TotalSeq-C barcoded MART-1 tetramers. Bulk expanded populations were analyzed using the 10x Genomics Single Cell 5′ Gene Expression and V(D)J workflow. **(a)** Harmonized UMAPs showing the method of expansion, distribution of CD4^+^ and CD8^+^ T cells, and MART-1 specificity, defined by staining with two independent barcoded MART-1 tetramers. Unsupervised clustering of the MART-1-specific population identified three distinct clusters. **(b)** Selection of differentially expressed genes by MART-1 specific T cells expanded with APC-ms and mo-DC by Bonferroni-corrected Wilcoxon rank-sum test. Average Z-scored expression across donors is shown. **(c)** Harmonized UMAP showing TCR clonotype expansion. Highly expanded clonotypes overlapped with the MART-1-specific cluster. **(d)** Top 100 most expanded MART-1-specific clonotypes expanded by APC-ms and mo-DCs, ranked by frequency of CD8^+^ T cells. The area under the curve (AUC) was calculated for each donor, and paired t tests were used to compare mean AUC values. **(e)** Expansion of clonotypes shared between APC-ms and mo-DC conditions for each donor. **(f)** Comparison of mean cell counts of shared clonotypes between APC-ms and mo-DCs. **(g)** TCR clonality of MART-1-specific clonotypes expanded with APC-ms and mo-DCs, calculated as the inverse of normalized Shannon diversity. **(h)** Schematic of the cohort comprising eight treatment-naive, MART-1^+^ melanoma patients. T cells isolated from peripheral blood were expanded for 12 days using the optimized APC-ms formulation described in Figure 1. **(i)** Representative MART-1-specific dextramer staining after 12 days of expansion. **(j)** Violin plot representing the distribution of MART-1-specific expansion (dextramer^+^ cells among CD8^+^ T cells) across 8 melanoma patients and 50 healthy donors. Solid line represents the median; dotted lines represent the 25^th^ and 75^th^ quartiles. **(k)** Schematic of patient Mel9 with genomically characterized melanoma expressing MART-1 and the personal neoantigen DCAKD p.S199F. **(l)** Representative antigen-specific dextramer staining after 12 days of expansion. **(m)** Functional validation of expanded antigen-specific T cells. Normalized CD137 expression was measured after 20 hours of coculture with autologous peptide-pulsed PBMCs.

Paired TCR sequencing showed that both platforms drove polyclonal expansion of MART-1-specific T cells, with clonotype sizes ranging from singletons to clones containing more than 400 captured cells (Fig. 2c). APC-ms expansion recovered a broader repertoire of unique MART-1-specific clonotypes and yielded a significantly greater proportion of expanded clones (clone size ≥ 1) within the MART-1-specific repertoire than mo-DCs (0.3223 vs. 0.1416, respectively; p = 0.031)(Extended Data Fig. 8b). The most abundant MART-1-specific clonotypes expanded by APC-ms constituted a larger proportion of the CD8^+^ T cell repertoire than those expanded by mo-DCs, however this difference was not statistically significant (Fig. 2d). Among clonotypes recovered by both platforms within the same donor, APC-ms drove greater expansion of individual clones than mo-DCs (Extended Data Fig. 8a, Fig. 2e,f). Moreover, clonality (the inverse of Normalized Shannon Entropy) of MART-1-specific TCRs expanded with APC-ms was greater than that of mo-DCs, indicating a less even repertoire with greater representation of highly expanded clonotypes (Fig. 2g). Together, these data show that APC-ms can support robust expansion of a wide repertoire of antigen-specific T cells from peripheral blood while maintaining a favorable combination of memory-associated and cytotoxic features.

### Recovery of MART-1-specific T cells from the blood of patients with melanoma

Having established performance in healthy donors, we asked whether the reference MART-1 formulation could recover circulating TRTs from patient material, in which precursor frequency and state may be shaped by prior antigen exposure and tumor-associated tolerance. T cells from peripheral blood of eight treatment-naive HLA-A*02:01+ melanoma patients were expanded for 12 days (Fig. 2h and Supplementary Table 3). APC-ms enriched MART-1-specific cells to a median of 13.7% of CD8+ cells (range 0.30 – 57.2%), with representative values shown in Fig. 2i. Expansion reached a threshold of greater than 5% in a greater proportion of patients (62.5%, 5 of 8 donors) than healthy donors (43%, 21 of 50 donors), demonstrating that the chosen reference formulation can enrich TRTs across several contexts (Fig. 2j).

### Parallel expansion of a shared antigen and a personal neoantigen

To test whether the platform could extend beyond a shared TAA and across different HLA alleles, we applied APC-ms to expand blood-derived T cells from a patient with melanoma (Mel9) that was previously genomic characterized to express candidate personal neoantigen arising from a DCAKD p.S199F mutation (Fig. 2k, Supplementary Table 4,5)^6,31,32^. The corresponding predicted 10-mer neoepitope (RFLEYLPLRF) was HLA-A*24:02-restricted. Circulating T cell responses against the mutant peptide were previously not detected *ex vivo* nor after a round of pre-stimulation with peptide^31^. However, functional testing of TIL-derived TCRs identified 5 TCRs reactive against the mutant peptide^6^. This provided an independently validated personal neoantigen with a defined HLA restriction for testing APC-ms expansion of rare TRTs from blood. As the patient also was HLA-A*02:01^+^, we had the opportunity to expand neoantigen- and TAA-specific T cells in parallel.

T cells from Mel9 peripheral blood were expanded with APC-ms at 0.1, 0.5 and 1 mol% biotinylated lipid presenting pHLA for MART-1/HLA-A*02:01 or DCAKD p.S199F/ HLA-A*24:02. 0.5 mol% yielded a significantly greater enrichment of MART-1-specific T cells than 0.1 mol%, further validating the choice of reference formulation (Extended Data Fig. 9, Fig. 2l). DCAKD-specific enrichment was greatest at 1 mol%, reaching 7.45% of CD8^+^ T cells (Fig. 2l). This requirement for a higher density of stimulation for expansion of DCAKD-specific T cells may reflect a low precursor frequency relative to MART-1-specific T cells as would be expected for a neoantigen-specific T cell^9^. Both antigen-specific populations upregulated CD137 in response to cognate peptide-pulsed autologous PBMCs (Fig. 2m), validating functional recognition of the peptide. Thus, the modularity of APC-ms enabled the parallel expansion of multiple T cell specificities from a single blood draw.

## Discussion

Together, these findings highlight APC-ms as a modular, cell-free platform for recovering and expanding functional antigen-specific T cells directly from peripheral blood. The use of the TAA, MART-1, as a model antigen in healthy donors allowed us to first screen stimulatory ligands presented on APC-ms to establish an optimized formulation for antigen-specific T cell expansion. In direct comparisons across multiple healthy donors, APC-ms outperformed conventional mo-DC cultures by inducing greater MART-1-specific enrichment and fold expansion while preserving polyfunctionality against a cognate target cell line. APC-ms also preserved an early differentiation phenotype (increased proportion of the CD45RA^+^CD62L, T_SCM_ compartment) and increased functional avidity against cognate peptide. Using the same reference formulation, APC-ms was used for expansion of MART-1-specific T cells from fifty healthy donors and nine melanoma patients. Furthermore, density titration enabled APC-ms to recover neoantigen-specific TRTs from peripheral blood of a patient with previously undetectable responses to the mutant peptide.

The success of APC-ms likely reflects the tunability of the material and its capacity to support T cell interactions in a physiological manner. In culture, the high aspect-ratio rods spontaneously assemble into three-dimensional scaffolds and elicit extensive T cell-material interactions, increasing the probability of a true antigen-specific T cell-material engagement ^22^. APC-ms also combines lateral ligand mobility with independent control of antigenic and co-stimulatory signals within a single, cell-free platform, differentiating it from alternative aAPC platforms based on polymeric scaffolds such as Xynapse. The ability to load defined molar amounts of ligands is a key advantage over cellular APCs like mo-DCs which may present antigen sub-optimally. This tunability enabled enrichment of T cells recognizing the personal neoantigen DCAKD and highlights ligand density as an important determinant of antigen-specific expansion. Finally, the defined stimulation provided by APC-ms may reduce the need for repeated rounds of stimulation that are common in traditional mo-DC-based protocols, which can require several weeks of ex vivo manipulation that may impair T cell performance. Here, APC-ms expanded TRTs following a single 12-day stimulation.

While these results underscore the potential of APC-ms as a tunable platform for antigen-specific T cell expansion, further studies are needed to characterize design rules for expansion and enable broader translational applicability. In particular, future work should determine how APC-ms stimulation variables influence antigen-specific activation. These effects are likely context dependent, reflecting features such as the T cell starting phenotype, precursor frequency, and pHLA-TCR avidity. As this study focused primarily on MART-1-specific expansion, validation across additional patient samples, HLA types, and antigen classes will be important. While mo-DCs are considered the gold-standard for antigen-specific expansion, emerging clinical expansion systems such as NEO-STIM, an optimized workflow for priming of neoantigen-specific T cells using patient-derived APCs, will also be informative benchmarks for establishing performance.^33^ Furthermore, additional characterization of APC-ms-expanded T cells will also be necessary to define the phenotypic, functional, and TCR-intrinsic properties selectively induced by the platform and their implications for therapeutic use.

APC-ms-mediated recovery of TRTs from the peripheral blood provides opportunities for both TIL therapy and therapeutic TCR discovery. Due to their rarity, discovery of peripheral blood TRTs has been limited and historically required extensive biomarker characterization and pre-enricment^8,11^. Here, we provide proof-of-principle that APC-ms can expand previously undetectable neoantigen-specific TRTs within 12 days. Moreover, in the MART-1 model system, expanded TRTs retained phenotypic and functional features that may be advantageous for adoptive cell therapy, including an early differentiation phenotype, polyfunctionality, and increased functional avidity,^9,34,35^. Collectively, these findings support APC-ms as a promising platform for both the discovery of tumor-reactive TCRs and personalized manufacturing of antigen-specific T cell products.

## Methods

### Human subjects and samples

Healthy donor peripheral blood mononuclear cells (PBMCs) were obtained from Brigham and Women’s Hospital as-de-identified apheresis collars. Patient PBMCs were collected on protocols approved by the Dana-Farber/Harvard Cancer Center Institutional Review Board, and all participants provided written informed consent in accordance with the Declaration of Helsinki. Samples from patient Mel9 were collected on NCT01970358^31^. PBMCs were isolated by density-gradient centrifugation and cryopreserved in 10% dimethyl sulfoxide in fetal bovine serum. Clinical characteristics are provided in Supplementary Tables 3 and 5.

### Cell Lines

Human melanoma cell lines SK-MEL-5 (HTB-70) and A375 (CRL-1619) and the human T2 lymphoplastoid cell line (CRL-1992) were obtained from the American Type Culture Collection (ATCC). Adherent lines SK-MEL-5 and A375 were maintained in Dulbecco’s Modified Eagle’s Medium (DMEM, ATCC 30-2002) supplemented with 10% fetal bovine serum (FBS) and 1% (P/S). When cells reached 80% confluence, they were passaged with 0.25% trypsin, 0.53 mM EDTA solution. T2 cells were maintained in ATCC-formulated Iscove’s Modified Dulbecco’s Medium (IMDM, ATCC 30-2005) supplemented with 20% FBS and 1% P/S. Medium was renewed every 2 days.

### APC-ms fabrication

Mesoporous silica rods were synthesized by a sol–gel route and characterized as described^22^. Representative dimensional analysis is shown in Extended Data Fig. 1. APC-ms was prepared as previously described with the following modifications^22,23^. Rods were coated with a supported lipid bilayer formed from extruded POPC liposomes containing 0.1, 0.5 or 1 mol% biotinylated lipid. Streptavidin was subsequently added at 35% theoretical molar saturation of the biotinylated lipid in the formulation and incubated in block solution (0.25% bovine serum albumin [BSA] in PBS, w/v) for 10 minutes at room temperature. Biotinylated peptide-HLA and biotinylated co-stimulatory molecules (Biolegend, αCD28 [CD28.2] Cat. No. 302904, αCD137 [4B4-1] Cat. No. 309806 or αICOS [C398.4A] Cat. No. 313505) were each added at 50% molar saturation of the added streptavidin, obtaining a 1:1 molar ratio. APC-ms was used immediately for T cell expansion experiments. Cytokines were supplied in the culture medium and were not loaded into the mesoporous core in this study.

### Peptides and peptide-HLA multimers

Biotinylated MART-1/HLA-A*02:01, DCAKD/HLA-A*24:02, and control peptide-HLA reagents were obtained from the NIH Tetramer Core Facility. Commercial MART-1 dextramers were used for flow cytometry (Immudex, WB02162) Barcoded tetramers were generated with five sequential additions of TotalSeq-C fluorescent streptavidin to the pHLA monomer. Peptides were synthesized by GenScript at greater than 95% purity. Antigen sequences and HLA restrictions are listed in Supplementary Table 4.

### Generation and characterization of mo-DCs

CD14^+^ monocytes were isolated from PBMCs of the same donors using CD14 MicroBeads (Miltenyi Biotec) and differentiated in Mo-DC Differentiation Medium (Miltenyi Biotec, cat. no. 130-094-812), followed by maturation in Mo-DC Maturation Medium according to the manufacturer’s instructions (Extended Data Fig. 4a–d). Maturation was verified by flow cytometry for canonical dendritic cell markers (CCR7, CD80, CD86, DC-SIGN, CD25, HLA class I, HLA class II) in all six donors. Mature mo-DCs were pulsed with 5 µg/ml of peptide and co-cultured with T cells from the same donor at a T cell:mo-DC ratio of 10:1. For benchmarking experiments, cultures received only a single stimulation with mo-DCs.

### Antigen-specific T cell expansion

T cells were isolated from PBMCs by immunomagnetic negative selection (Pan T isolation kit, Miltenyi) and co-cultured with APC-ms or peptide-pulsed mo-DCs in X-VIVO-15 medium (Lonza, 04-418Q) supplemented with 5% human serum. Cultures were seeded at 2 x 10^6^ cells per well in a 48 well plate. Between two and four technical replicate wells were established per condition as indicated and according to sample availability. Each donor received a single stimulation with APC-ms on day 0. IL-21 (30 ng/ml) was present from day 0 to day 3, followed by IL-7 and IL-15 (each 5 ng/ml) from day 3 onward. IL-2 was used at 100 IU/ml where indicated. Medium was exchanged every 3 days. Antigen-specific enrichment and surface phenotype were assessed on day 12. Cytokine profiling and CD137 dose-response assays were performed on day 18 to reduce the activation background present early post-stimulation. Xynapse reagents (Immudex) were prepared according to the manufacturer’s instructions and used in parallel cultures under the same soluble cytokine schedule.

In tuning experiments to expand MART-1-speciifc T cels from blood, donor-level differences across densities were not statistically significant unless otherwise indicated in Extended Data Fig. 3. 0.5 mol% lipid biotinylation was selected for subsequent experiments as it balanced robust MART-1-specific expansion while reducing peptide–HLA reagent requirements and associated costs. αCD28 co-stimulation achieved maximal MART-1-specific expansion that was not improved with the addition of alternative co-stimulatory molecules. IL-21 supplementation during the first three days of stimulation enriched for MART-1-specific T cells within the T_SCM_ compartment. Low-dose IL-7 and IL-15 were chosen for supplementation during extended culture based on their established role in supporting development of memory T cells.^29^

### Flow cytometry, multimer staining and phenotyping

Cells were stained with tetramer or dextramer for 15 minutes at room temperature before the addition of surface antibodies and viability stain. The gating strategy is shown in Extended Data Fig. 2. Data were acquired on a BD LSRFortessa and analyzed in FlowJo v10^10^.

### Functional avidity and activation-induced marker assays

For the MART-1 platform comparison, T2 cells were pulsed with the indicated concentrations of cognate peptide and co-cultured with expanded T cells. For the Mel9 antigen-specific assays, EBV-lymphoblastoid cell lines (EBV-LCLs, provided by Oliveira et a.^6^) were pulsed with peptide for 2 hours and used as targets. CD137 upregulation on antigen-specific CD8+ T cells was measured after 20 hours^36^. Background from unstimulated targets was subtracted, and half-maximal effective concentrations were estimated by four-parameter logistic regression in GraphPad Prism version 11.0.1.

### Intracellular cytokine staining and degranulation

Expanded T cells were challenged with SK-MEL-5 (MART-1^+^HLA-A*02:01^+^) or A375 (HLA-A*02:01^+,^ MART-1^-^) cells at an effector-to-target ratio of 10:1 in the presence of anti-CD107α. Brefeldin A was added after 4 hours. After 6 hours, cells were surface stained, fixed (Fixation Buffer, Biolegend, Cat. No. 420801), permeabilized (Intracellular Staining Permeabilization Wash Buffer (10X), Biolegend, Cat. No. 421002), and stained with IFNγ, TNFα, and IL-2 antibodies. Unstimulated cultures and phorbol myristate acetate (PMA) with ionomycin served as negative and positive controls respectively.

### Preparation of cells for transcriptomics

Day 12 cultures from three donors expanded in parallel with APC-ms or mo-DCs were stained with TotalSeq-C-barcoded antibodies and peptide–HLA tetramers assembled on barcoded streptavidin (Biolegend, listed in Supplementary Table 1). Samples expanded from APC-ms and mo-DC from the same donor were stained with TotalSeq-C Hashtag Antibodies before pooling and running on the same sequencing lane. Two independently barcoded tetramers were prepared per specificity, and only cells positive for both MART-1 barcodes were assigned as antigen specific. Barcoded tetramers presenting an Epstein–Barr virus epitope and an unrelated HLA-A*02:01-restricted neoantigen were included to quantify nonspecific multimer binding (Extended Data Fig. 6). Libraries for 5′ gene expression, V(D)J, and feature barcoding were prepared using the 10x Chromium Single Cell 5’ and V(D)J Enrichment kit and sequenced on the NovaSeq 6000 (S4) system.

### Processing of single-cell RNA sequencing data

Reads were aligned and quantified with cellranger-8.0.1 (10x Genomics) with the GRCh38-2020-A reference transcriptome. CITE-seq data were processed jointly with the corresponding gene expression (GEX) data. Antibody-derived tag (ADT) counts and GEX Feature Barcode analysis were contained within the same count matrix for downstream analysis. scRNAseq data was processed using standard pipelines in Seurat V5^36^. Cells that contained greater than 10% mitochondrial RNA content, unique feature counts less than 250 or greater than 7000, and less than 500 unique molecular identifiers (UMIs) were excluded from the analysis. This yielded 49,526 cells across three donors expanded with APC-ms and mo-DCs in parallel. The data was then normalized using Seurat NormalizeData function (LogNormalize method) and CITE-seq data using the center log-ratio (CLR) function. ScaleData, FindVariableFeatures and RunPCA functions in Seurat were used for data integration and dimensionality reduction. RunHarmony was used for batch correction. The HTOdemux function was used to deconvolute hashtag labels and determine cell identity. Cells that reached a user-defined threshold for CLR-normalized expression of both MART-1 tetramers while remaining below threshold for the two negative control tetramers were labeled as MART-1-specific. This process labeled 32,587 cells from APC-ms (3,293 of which were MART-1-specific) and 16,436 cells from mo-DCs (1,216 of which were MART-1-specific). MART-1-specific T cells were resolved into three clusters by the FindNeighbors and FindClusters functions in Seurat. Differential expression analysis based on the non-parameteric Wilcoxon rank sum test was implemented between all MART-1-specific cells comparing APC-ms and mo-DCs using FindMarkers(). Differentially expressed genes are those that had Bonferroni-adjusted p values (p_val_adj) less than or equal to 0.05. The full list of differentially expressed genes is shown in Supplementary Table 2. The propeller function implemented within the speckle package was used to generate variance-stabilized normalized proportions of MART-1 specific cells from APC-ms and mo-DC expanded samples belonging to each MART-1-specific cluster.

### Processing of single-cell TCR sequencing data

Paired V(D)J sequencing data were processed using Cell Ranger v8.0.1 (10x Genomics) with the vdj_GRCh38_alts_ensembl-5.0.0 V(D)J reference. Individual TCRs clones were grouped and filtered as previously described by Oliveira et al.^6^ Clonotype families were formed on the basis of shared TCRα or TCRβ sequences, allowing for single amino acid substitutions within the CDR3 domain. The frequency of each clonotype family was calculated separately for each donor and expansion method and considered a single clonotype. Repertoire clonality was calculated as the inverse of normalized Shannon entropy^37^.

### Statistics and reproducibility

Two-sided paired t tests were used for donor-matched platform comparisons where indicated in the figure legends. For Fig. 2e, the area under each donor’s rank-frequency curve was calculated and donor-level values were compared with a paired t test. No adjustment for multiple comparisons was applied to donor-level tests unless stated. Differential expression between MART-1-specific cells expanded with APC*-*ms and mo-DCs was assessed using the Bonferroni-corrected Wilcoxon rank-sum test at the cell level. Donors and patients were treated as independent biological replicates. P < 0.05 was considered significant unless otherwise noted. Technical replicate wells from the same donor were not treated as independent biological replicates. Summary bars show mean ± SEM. unless otherwise indicated, with individual biological replicates displayed where applicable. No statistical method was used to predetermine sample size, and no samples were excluded.

## Data availability

Single-cell gene-expression, V(D)J and CITE-seq data will be deposited in the Gene Expression Omnibus repository.

## Author contributions

D.J.M., C.J.W. and N.C. conceived and designed the study. Y.B., D.Z., J.D. and V.M.D. developed, fabricated and characterized the antigen presenting cell mimetic scaffolds. N.C., J.D. and H.J. performed T cell expansion, flow cytometry and functional studies. J.D. performed computational analyses. M.M. provided treatment-naive melanoma samples. G.O. provided patient samples, cell lines and peptides relevant to Mel9. D.N. contributed to statistical design and analysis. P.A.O. and J.R. contributed to data discussion and interpretation. D.J.M. and C.J.W. supervised the study. J.D. and Y.B. wrote the manuscript with input from all authors. All authors reviewed and approved the final manuscript.

## Acknowledgements

We thank the NIH Tetramer Core Facility for production of peptide-HLA reagents and for technical guidance. This research was supported by the National Cancer Institute of the National Institutes of Health under Award Number U54CA244726. JD is supported by the National Science Foundation Graduate Research Fellowship. YB was supported by the National Cancer Institute under training grant T32CA136432. NC is supported by NMDP/Be the Match – Amy Strelzer Manasevit Research Program, Deutsche Knochenmarkspendedatei (John Hansen Research Grant) and the Claudia Adams Barr Program for Innovative Cancer Research. NC is also a Scholar of the American Society of Hematology, and of Blood Cancer United, formerly The Leukemia & Lymphoma Society (Career Development Program – Special Fellow). DJM is the Pinkas Family Professor of Bioengineering at Harvard University, and his work is supported, in part, by the Wyss Institute. CJW is the Lavine Family Chair for Preventative Cancer Therapies, and a member of the Parker Institute for Cancer Immunotherapy at Dana-Farber Cancer Institute. Her work is supported, in part, by the Parker Institute for Cancer Immunotherapy.

