## Supplemental Data for "Programmable antigen-presenting scaffolds expand rare tumor-antigen-specific T cells from peripheral blood"

### Extended Data

**Extended Data Fig. 1: Mesoporous silica rod characterization**. Distribution of (a) length and (b) width of mesoporous silica rods by image-based annotation.

**Extended Data Fig. 2: Gating strategy for T cell characterization.** Gating strategy for dextramer staining of APC-ms expanded T cells (Fig. 1b) and phenotypic characterization of T cells expanded with APC-ms or mo-DCs (Fig 1d-h). Expanded T cells were analyzed by flow cytometry on Day 12.

**Extended Data Fig. 3: Donor-level expansion outcomes during APC-ms tuning.** MART-1-specific enrichment (dextramer$^+$ cells among CD8$^+$ T cells, top, n=5 donors), fold expansion (middle, n = 3 donors), and percentage of MART-1-specific of $T_{SCM}$ cells (bottom, n = 4 donors) across the tuning conditions shown in Fig. 1b. Statistical significance was assessed using paired t tests; nonsignificant comparisons are not shown ($p < 0.05$). While 1% presentation density induced the greatest enrichment and fold expansion across donors, these differences were not statistically significant. The presentation density of 0.5% was chosen for the final formulation to balance cost and expansion performance. Anti-CD28 alone was sufficient to achieve maximal enrichment of dextramer$^+$ T cells. Finally, sequential supplementation with IL-21 followed by IL-7 and IL-15 on day 3 produced the greatest enrichment while maximizing the proportion of $T_{SCM}$ cells.

**Extended Data Fig. 4: Generation and characterization of mo-DCs and expanded T cells.** **(a)** Workflow for mo-DC generation using the Miltenyi mo-DC Generation Toolbox. **(b)** Numbers of cells recovered during mo-DC generation workflow. **(c)** Representative histograms showing expression of canonical DC markers. **(d)** Heatmap showing mean expression of DC characterization markers across 6 donors. **(e)** Bubble plot of expansion across 6 donors with APC-ms or mo-DCs. **(f)** Expression of activation-induced and inhibitory markers by flow cytometry across 6 donors expanded with APC-ms or mo-DCs. Data correspond to the markers shown in the radar plot in Figure 1h.

**Extended Data Fig. 5: Comparison of APC-ms expansion with commercial artificial APC platform: Xynapse (Immudex).** Xynapse is a dextran-scaffold-based expansion technology presenting pHLA and co-stimulatory signals. T cells from 3 healthy donors were expanded with either APC-ms or Xynapse according to the manufacturer's instructions. **(a)** Representative dextramer staining of T cells expanded with APC-ms or Xynapse. **(b)** MART-1-specific CD8$^+$ T cell enrichment and fold-expansion across 3 healthy donors. Error bars represent the mean ± SEM and p values were calculated using paired t tests (*: $p \le 0.05$, **: $p \le 0.01$).

**Extended Data Fig. 6**: **Expression of CITE-seq antibodies and tetramers.** Two tetramers per specificity were prepared with TotalSeq-C-barcoded streptavidin. Cells stained by both MART-1 tetramers were classified as MART-1–specific. Negative-control tetramers targeting an HLA-A*02:01-restricted EBV BMLF1 epitope (GLCTLVAML) and an HLA-A*02:01-restricted neoantigen, TMEM214 p.S605F (QLPDFVNQL), were included to assess nonspecific tetramer staining.

**Extended Data Fig. 7: Distribution of APC-ms and mo-DC-expanded cells among MART-1-specific clusters.** MART-1-specific subcluster proportions were calculated within each donor-condition sample (relative to total MART-1$^+$ cells) and compared between APC-ms and mo-DC conditions using propeller (speckle package) which yields variance-stabilized, donor-level differential proportion analysis. No comparison reached levels of statistical significance.

**Extended Data Fig. 8**: **Supplemental TCR sequencing analyses.** (a). Upset plot showing shared and unique TCR clonotypes across conditions and donors. (b) Pie chart showing the clonal composition of MART-1-specific CD8$^+$ cells across donors expanded with APC-ms or mo-DCs. Each color is an individual clonotype. The total number of cells recovered, and number of unique clonotypes recovered are annotated. Singlets are pooled and shown in gray.

**Extended Data Fig. 9**: **Peptide-HLA density optimization in patient Mel9.** MART-1- and DCAKD p.S199F-specific CD8$^+$ T cell enrichment across APC-ms formulations containing 0.1%, 0.5%, or 1% biotinylated lipid displaying MART-1 or DCAKD p.S199F pHLA. Symbols represent technical replicates from the same donor.

**Tables**

**Supplementary Table 1**: TotalSeq-C reagents for CITEseq

**Supplementary Table 2:** Differentially expressed genes by MART-1-specific T cells expanded with APC-ms or mo-DCs

**Supplementary Table 3**: Clinical characteristics of the eight treatment-naive melanoma patients.

| Pt ID | Age | Sex | Primary Site | Stage |
|---|---|---|---|---|
| Mel1 | 60 | M | Back | T2a, locally advanced |
| Mel2 | 66 | M | Anus | T2, locally advanced |
| Mel3 | 82 | M | metastatic | metastatic |
| Mel4 | 44 | F | scalp | local |
| Mel5 | 39 | F | upper arm | pT3bN0, local |
| Mel6 | 83 | M | | |
| Mel7 | 70 | F | | metastatic |
| Mel8 | 79 | M | Back | T1aN3b, metastatic |

**Supplementary Table 4**: Peptides used for stimulation

| Antigen | Epitope | Sequence (mutant / wild type) | HLA restriction | Basis of restriction |
|---|---|---|---|---|
| MART-1 (MLANA) | MART-1 26–35 (A27L) | ELAGIGILTV | HLA-A*02:01 | Anchor-modified analog of the immunodominant Melan-A/MART-1 26–35 epitope[26,27] |
| DCAKD p.S199F | DCAKD 198–207 | RFLEYLPLRF / RSLEYLPLRF | HLA-A*24:02 | Experimentally deconvoluted by monoallelic HLA screening of five autologous TIL-derived TCRs[6] |

**Supplementary Table 5**: Clinical characteristics, HLA genotype and cross-study identifiers for patient Mel9.

| **Field** | **Value** |
|---|---|
| Identifiers | Mel9 (this study); Patient 1 (Ott *et al.*, Nature 2017)[31]; Pt-1 (Hu *et al.*, Nat Med 2021)[32]; Pt-A (Oliveira *et al.*, Nature 2021)[6] |
| Age / sex | 26 / male |
| Primary site | Back |
| Site of resected disease | Axillary lymph node metastasis |
| Stage | IIIC (T3bN3M0) |
| Previous treatment | Interferon-α |
| Treatment after surgery | Personal neoantigen peptide vaccination (NeoVax, NCT01970358) |
| Recurrence after NeoVax | Yes, 40 months after surgery, brain |
| Treatment after recurrence | Surgery, radiation, BRAF/MEK inhibition and immune checkpoint blockade |
| HLA-A | 02:01, 24:02 |
| HLA-B | 44:02, 15:01 |
| HLA-C | 05:01, 07:02 |
| Sample used here | Pre-vaccination PBMC |

a

b

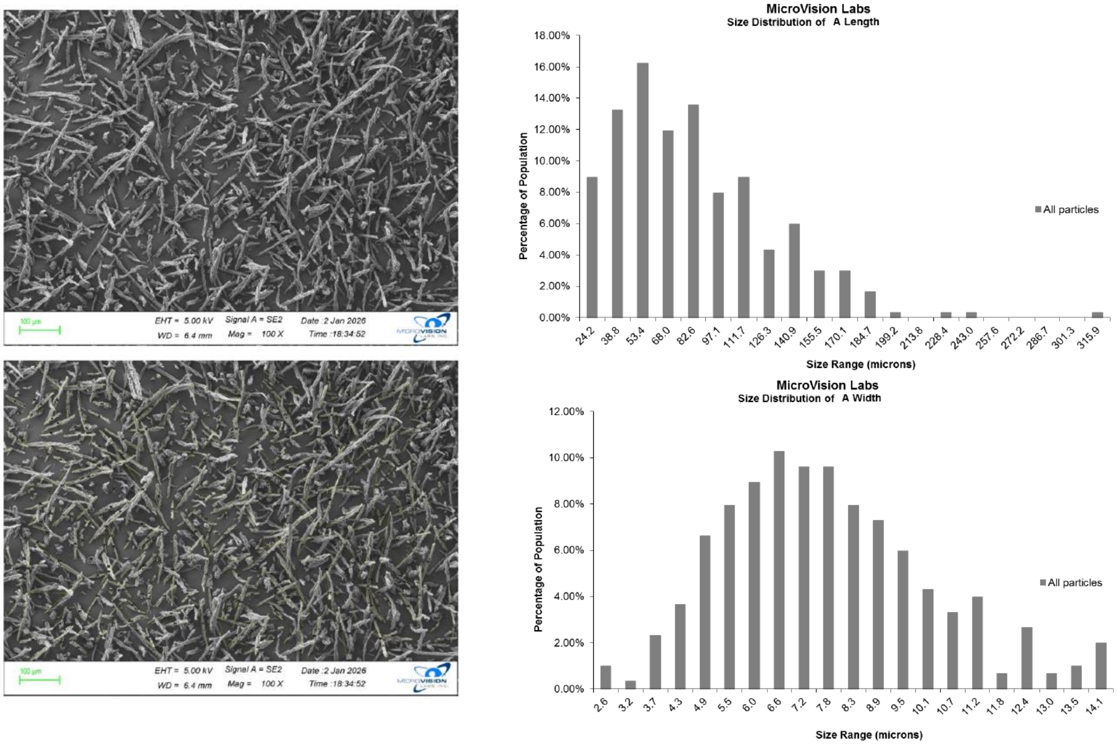


**S1 :MSR Characterization. Distributions of MSR (a) length and (b) width.**

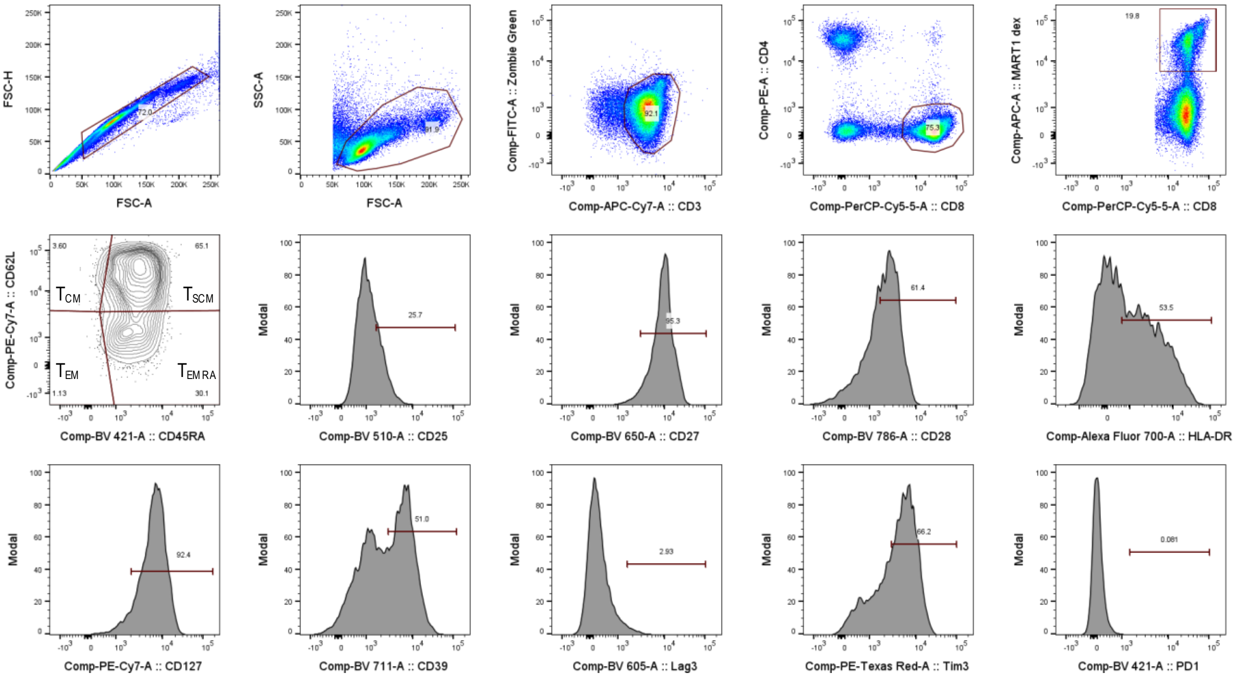


**S2: Gating strategy for T cell characterization.**

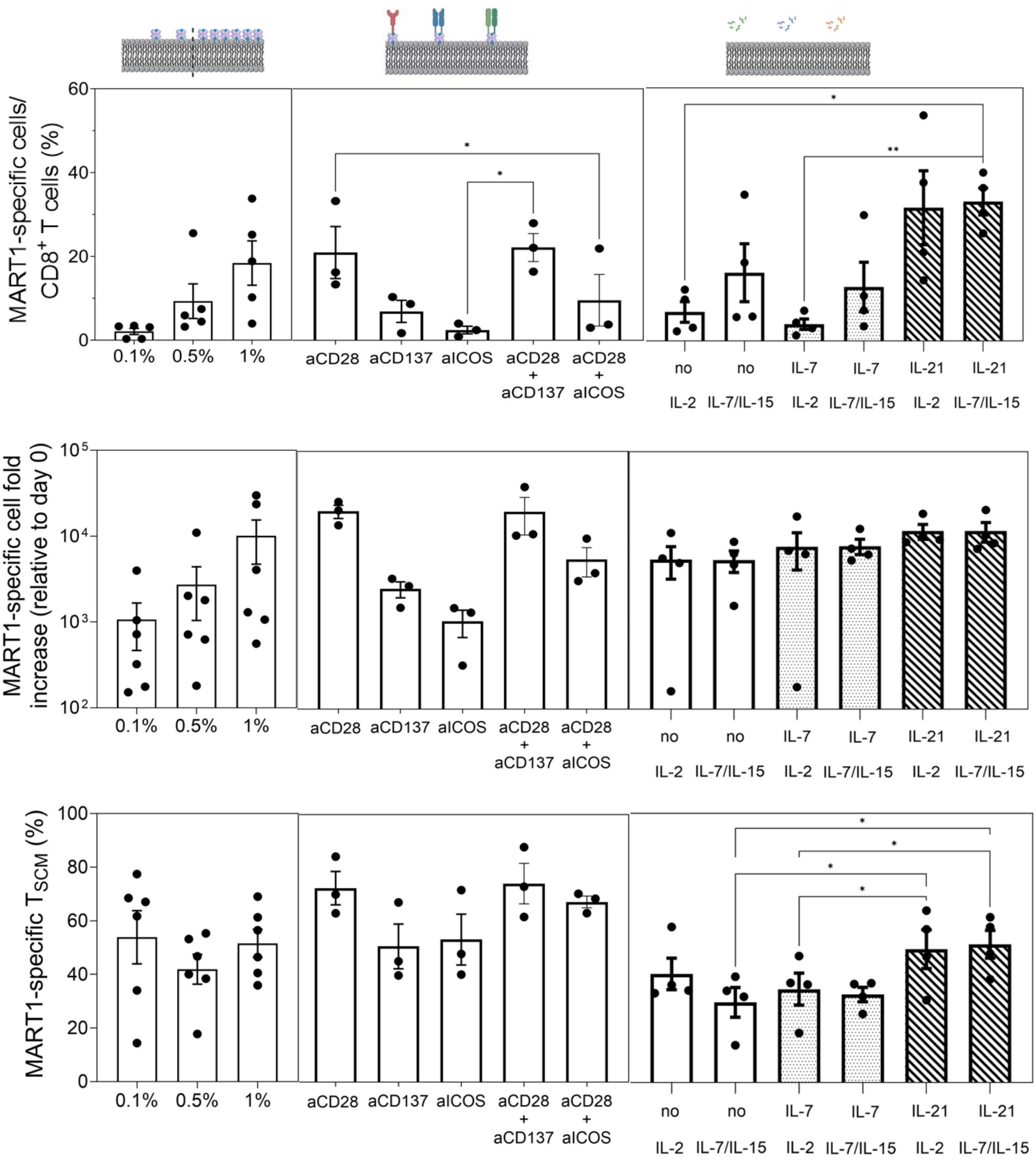


**S3 Donor-level expansion outcomes during APC-ms tuning.**

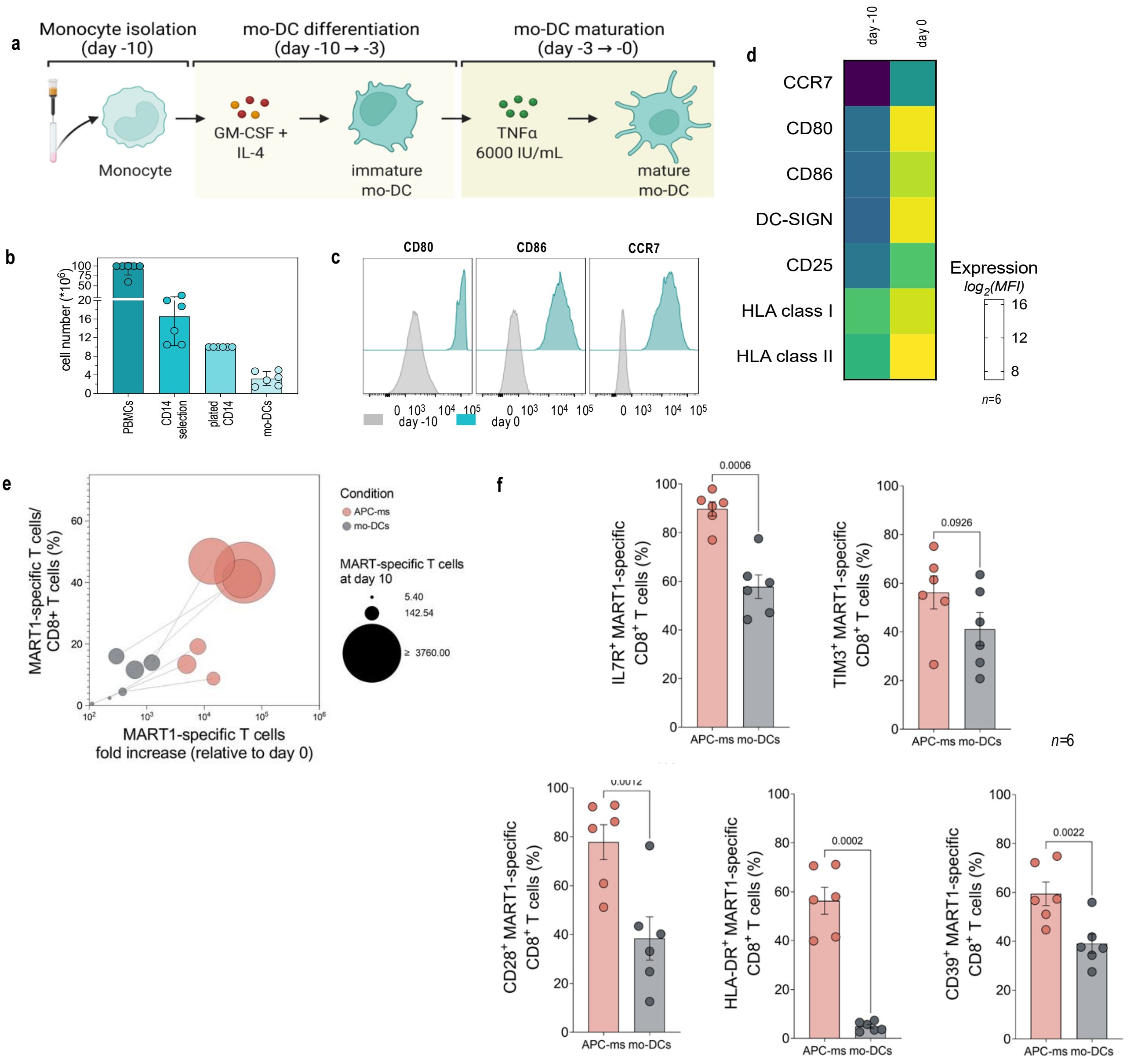


**S4. Generation and characterization of mo-DCs and expanded T cells.**

*n=3*

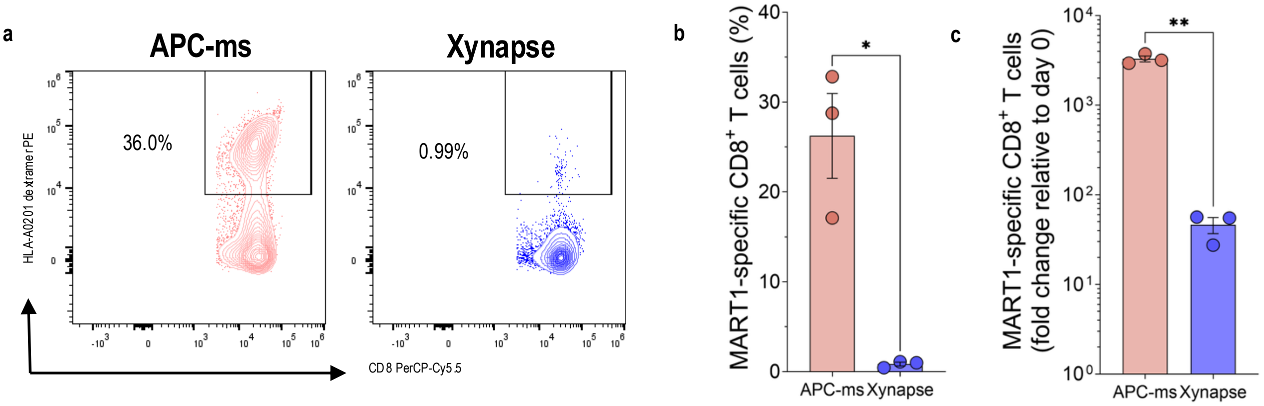


**S5. Comparison of APC-ms expansion with commercial artificial APC platform: Xynapse (Immudex).**

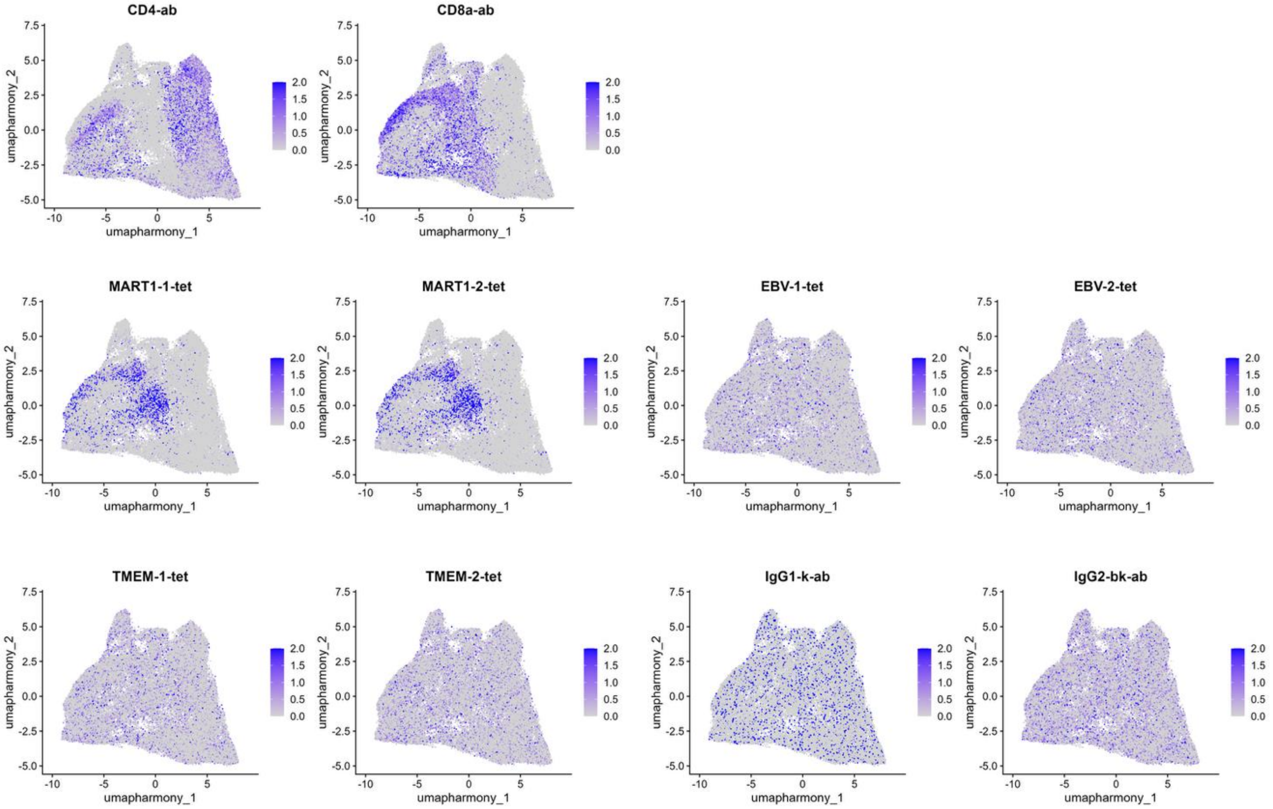


**S6. Extended Data Fig. 6: Expression of CITE-seq antibodies and tetramers.**

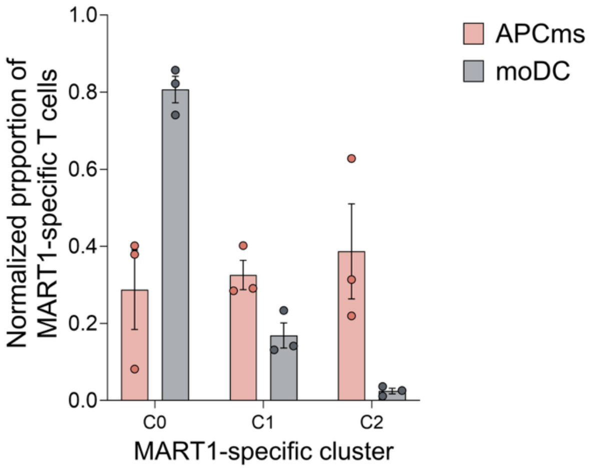


**S7. Distribution of APC-ms and mo-DC-expanded cells among MART-1-specific clusters**

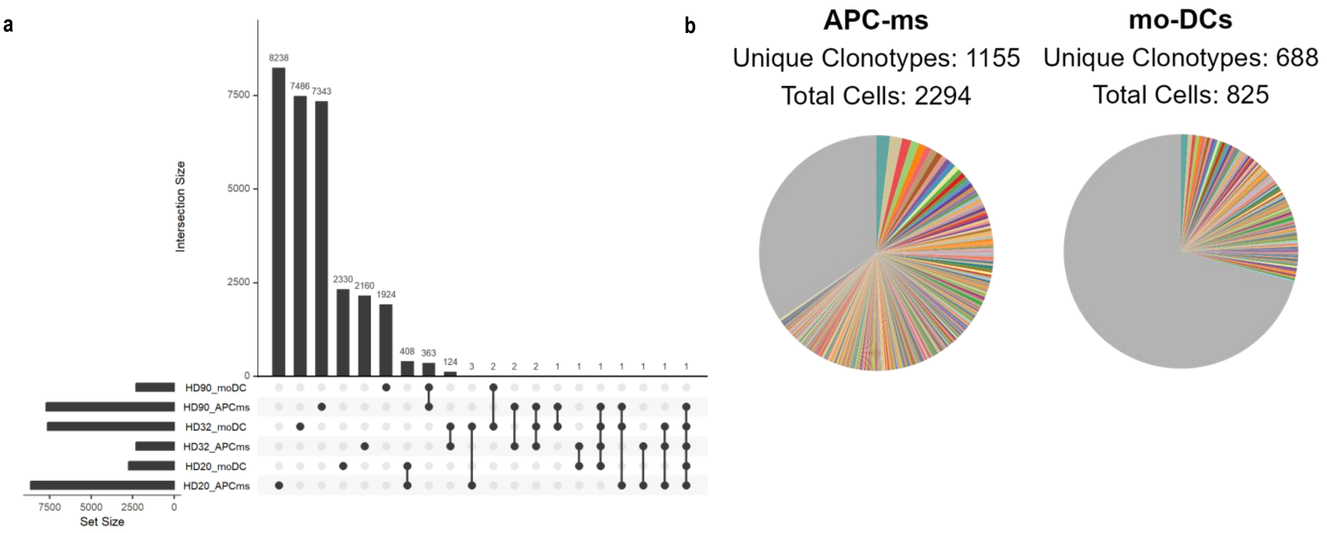


**S8. Supplemental TCR sequencing analyses.**

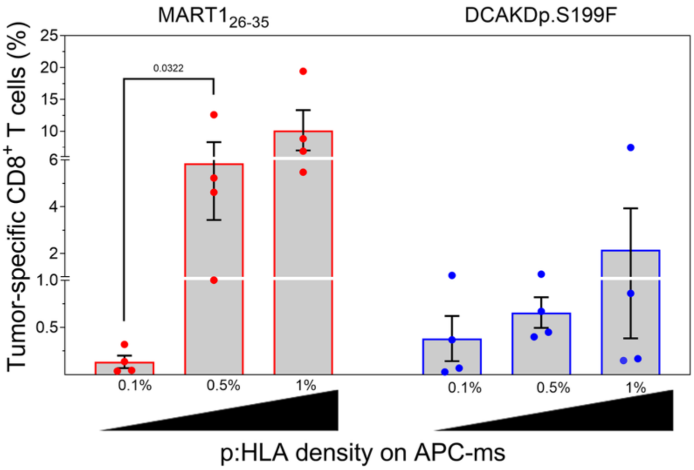


**S9. Peptide-HLA density optimization in patient Mel9.**
